# Lactucopicrin: A sesquiterpene lactone with anti-inflammatory activity modulates the crosstalk between NF-kB and AHR pathways

**DOI:** 10.1101/2025.08.02.667913

**Authors:** María Ángeles Ávila-Gálvez, Catarina J G Pinto, Carlos Rafael-Pita, Inês P. Silva, Aleksandra T. Janowska, Sérgio Marinho, Rory Saitch, Yilong Lian, Pakavarin Louphrasitthiphol, Jonas Protze, Gerd Krause, Pedro Moura-Alves, Cláudia Nunes Dos Santos

**Author notes:** Corresponding authors: Cláudia Nunes Dos Santos,; Pedro Moura-Alves.

## Abstract

Chronic inflammatory diseases, including inflammatory bowel diseases and cardiovascular diseases, share inflammation as a central pathological feature. The transcription factors NF-kB and AHR are key regulators of inflammatory responses, yet their interplay remains underexplored in the context of natural anti-inflammatory compounds. This study investigates the anti-inflammatory properties of sesquiterpene lactones derived from *Cichorium intybus* L. (chicory), focusing on their modulation of NF-kB and AHR signaling pathways. Using a TNFα-induced inflammation model in macrophages, endothelial, and intestinal epithelial cells, we identified lactucopicrin as a potent NF-kB antagonist. Notably, *in silico* docking and functional assays revealed lactucopicrin as a novel AHR modulator. Crucially, silencing AHR expression attenuated lactucopicrin-mediated NF-kB inhibition, uncovering a previously unrecognized AHR-NF-kB crosstalk mechanism. These findings not only position lactucopicrin as a dual-pathway modulator but also highlight the therapeutic potential of chicory-derived sesquiterpene lactones in treating inflammation-driven diseases and opening new avenues for leveraging natural products in targeted anti-inflammatory strategies.

## INTRODUCTION

Chronic diseases, such as inflammatory bowel diseases (IBD), cancer, cardiovascular diseases (CVDs), diabetes, and neurodegenerative diseases, are leading causes of death and disability worldwide^1–3^. Inflammation is a common hallmark across these conditions and is often exacerbated by environmental and lifestyle-related factors such as Western diets, stress, and smoking^1–3^. In particular, IBDs, including Crohn’s disease and ulcerative colitis, involving chronic intestinal inflammation are increasingly prevalent^4^ and impose significant costs on healthcare systems, with over 2 million Europeans affected^5^. So far, the current therapies are primarily chemical drugs and often fail for many patients^6^.

Gut inflammation can trigger an immune response, leading to the release of pro-inflammatory cytokines that reach circulation, affecting distant organs, including the vascular endothelium^7^. Remarkably, one of the hallmarks described in IBD patients is arterial stiffness due to chronic inflammation leading to endothelial dysfunction, promoting platelet aggregation and plaque formation^8^. Consequently, a connection between IBD and CVDs has been described^9,10^.

Among the key regulators of inflammation is the transcription factor nuclear factor kappa-light-chain-enhancer of activated B cells (NF-kB), a master regulator of inflammation and host defense against different insults, including microbial infections^11^. The NF-kB signaling pathway regulates gene expression of many cytokines and chemokines, as well as adhesion molecules, cell cycle regulators, and anti-apoptotic factors involved in different cellular, tissue, and organismal responses^12,13^. However, when deregulated, it is involved in various inflammatory processes such as IBD or endothelial dysfunction^14,15^. Another important regulator is the Aryl Hydrocarbon Receptor (AHR). This highly conserved ligand-dependent transcription factor regulates complex transcriptional programs in various cellular and tissue contexts, including during inflammatory responses^16^. Exogenous ligands include xenobiotics, microbial molecules, and metabolites, as well as diet-derived ligands like indigo, which have been described as ligands of AHR^16–19^. Notably, a recent review of over 100 microbial metabolites derived from dietary (poly)phenols identified eight as AHR modulators^20^. The ability of the AHR to bind a vast array of molecules, including bacterial signaling molecules, has been described, demonstrating that this binding is a crucial mechanism for sensing environmental cues and, therefore, host defense^17–19^.

Importantly, NF-kB is a key component that regulates AHR expression and induces AHR-dependent gene expression in immune cells^21^. Moreover, it has been described that inflammatory stimuli and cytokines that regulate NF-kB induce AHR expression, disclosing the existing crosstalk between AHR and NF-kB signaling pathways^21^. Therefore, targeting the NF-kB and AHR signaling pathways represents an attractive approach to tackling inflammatory processes. For instance, targeting the AHR has also been proposed for the treatment of cancer^22^, as well as for treating certain inflammatory skin conditions, such as dermatitis and psoriasis^23,24^, and inflammatory gut conditions, including Crohn’s disease^22^. The natural compound and AHR agonist tapinarof has been approved as a topical treatment for plaque psoriasis^23^.

Natural products represent a rich source of structurally diverse bioactive molecules with therapeutic potential. In this context, sesquiterpene lactones (STLs), a class of phytochemicals abundant in *Cichorium intybus* L. (chicory), have shown promising anti-inflammatory activity^24,25^. In this regard, the most abundant STLs in chicory are lactucin (LC), 11β,13-dihydrolactucin (DHLC), lactucopicrin (LCP), and 11β,13-dihydrolactucopicrin (DHLCP)^26^. Despite their potent biological activities, a significant obstacle limiting their pharmacological use is their poor aqueous solubility and low oral bioavailability, which may necessitate higher doses and increase systemic toxicity^27,28^. Following oral consumption of chicory preparations, a human pharmacokinetic study confirmed only LC and DHLC were detected in circulation at low concentrations,^29^ while in vitro fermentation assays demonstrated the conversion of LCP and LC into lactucin-type compounds by the gut microbiota^30^. These findings indicate that chicory STLs undergo extensive presystemic metabolism involving microbial and phase II conjugation processes, resulting in very low plasma levels. Therefore, mechanistic in vitro studies are essential to elucidate their biological targets and mechanisms of action, which may operate locally or at low systemic concentrations.

To overcome these challenges, recent efforts have focused on improving the delivery of STLs and related compounds through advanced drug delivery systems^31,32^. Nanoformulation approaches have been proposed as a promising strategy to enhance targeted delivery of hydrophobic natural products, thereby improving their therapeutic efficacy while minimizing off-target effects. For example, a study employing nanocarriers such as polylactide-co-glycolide nanoparticles or functionalized nanographene has successfully enhanced the delivery and bioactivity of parthenolide in preclinical models^31^.

Albeit the inhibition of the NF-kB pathway by numerous STLs has been well-characterized^25^, the ability of these chicory STLs to modulate the crosstalk between the AHR and NF-kB pathways remains unexplored, so far. In this study, we have evaluated the main STLs from *Cichorium intybus* L. (chicory) to identify inflammatory modulators, namely those capable of impacting the NF-kB signaling pathway. Taking advantage of a TNFα-induced inflammation model in diverse cell types, including macrophages and endothelial cells, we examined the impact of STLs on the modulation of NF-kB activity through gene and protein expression analysis, as well as by utilizing NF-kB signaling reporter systems. Among the STLs tested herein, our results reveal LCP as a major NF-kB antagonist. Furthermore, *in silico* docking analysis predicted that LCP binds and modulates the AHR, a prediction that was further confirmed by gene and protein expression, AHR signaling reporter systems, and enzymatic activity analysis in macrophages, endothelial cells, and intestinal epithelial cells. Finally, silencing AHR expression in endothelial cells attenuated the LCP antagonism observed in a model of TNFα-induced NF-kB activation, unveiling its impact on the AHR-NF-kB signaling crosstalk.

## RESULTS AND DISCUSSION

### Anti-inflammatory potential of LCP modulating NF-kB activity

Lactucin (LC), 11β,13-dihydrolactucin (DHLC), lactucopicrin (LCP), and 11β,13-dihydrolactucopicrin (DHLCP) list among the most abundant STLs found in chicory roots^33,34^, henceforth selected as candidates to be tested (**Figure 1A**). To evaluate the anti-inflammatory potential of each of the chosen STLs, we first assessed their capacity to modulate NF-kB activity using a well-established cell-based luciferase reporter system^18,19,35–37^.

**Figure 1.**
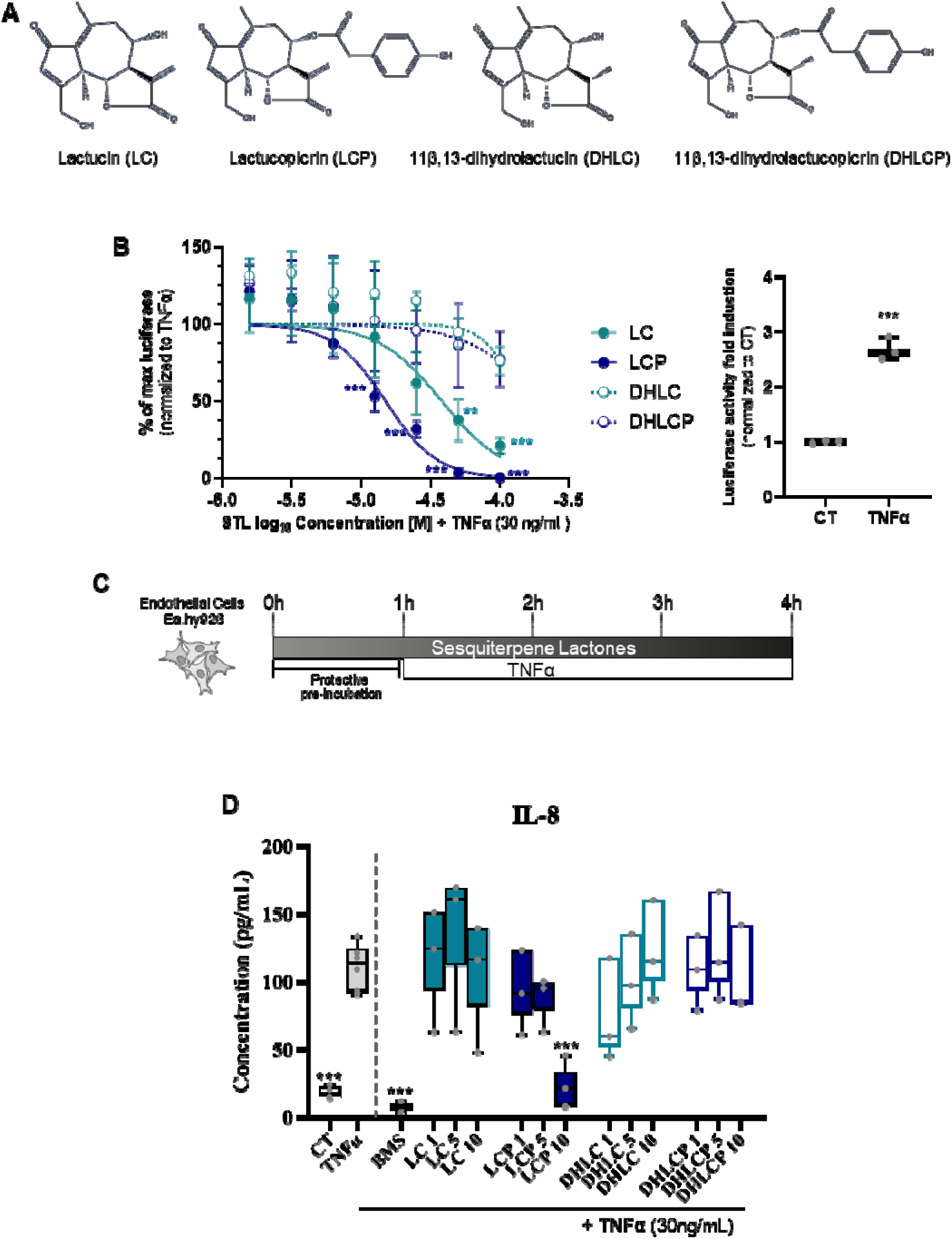
Screening of the anti-inflammatory effects of the four main chicory STLs. **A**) Chemical structures of the four STLs used. **B**) THP-1 (human monocytic) NF-kB luciferase reporter cells readout upon 4h exposure to 0.5% DMSO (CT) or TNFα (30 ng/mL) (left panel) or to different concentrations of lactucin (LC), lactucopicrin (LCP), dihydrolactucin (DHLC), or dihydrolactucopicrin (DHLCP) in the presence of TNFα (30ng/mL, right panel). **C**) Experimental design of the endothelial inflammation model used to evaluate the prophylactic effects of DHLC, DHLCP, LC, or LCP. **D**) EA.hy926 endothelial cells IL-8 release upon exposure to CT, TNFα, or TNFα in the presence of different concentrations of the STLs or the NF-kB inhibitor BMS-345541 (BMS, 5 µM). Data shown as means + SD (n=3). Statistical significance assessed by one-way ANOVA with multiple comparisons to TNFα; \*\**p*<0.01; \*\*\**p*<0.001.

In brief, THP-1 NF-kB luciferase reporter macrophages were treated with the different STLs in the presence (**Figure 1B**, left panel) or absence (**Supporting information; Figure S1**) of TNFα (30 ng/mL; NF-kB signaling pathway activator-positive control (**Figure 1B**, right panel), and luminescence assayed as a readout of NF-kB activation. Contrary to TNFα (**Figure 1B**), exposure to the different STLs did not lead to NF-kB activation (**Supporting information; Figure S1**). However, a decrease in basal luminescence was observed upon stimulation of LCP from 25µM to 100 µM, and to a lesser extent of LC at 100µM (**Supporting information; Figure S1**). To further assess the capacity of the different STLs to block NF-kB activation, we performed co-stimulation experiments in the presence of TNFα. LC and LCP significantly reduced TNFα-induced NF-kB activation (**Figure 1B**), with the LCP effects being stronger (IC_50_ = 10.6 µM) compared to LC (IC_50_ = 33.9 µM). Led by the results obtained using the THP-1 NF-kB reporter system in macrophages as a first screening, we evaluated the effect of the STLs in an endothelial inflammation model upon stimulation of human endothelial cells (EA.hy926 ^38^) with TNFα^39^. Of note, the upregulation of IL-8 expression in endothelial cells upon NF-kB activation by TNFα has been described^39^. Therefore, we assessed IL-8 expression levels as a readout of TNFα-induced inflammation. In brief, following a prophylactic approach model (**Figure 1C**), we assessed IL-8 levels in EA.hy926 upon exposure to TNFα alone or after pre-incubation with the different STLs, using the highest concentration, which was a value close to the ICLL of LCP previously determined in the THP-1 screening. These concentrations were non-cytotoxic to endothelial cells (**Supporting information; Figure S2**). Additionally, we used the well-characterized NF-kB inhibitor BMS-345541 (BMS) as a positive control^40^ (**Figure 1D**). Exposure to TNFα (30 ng/mL) elicited IL-8 production, which was attenuated in the presence of the NF-kB inhibitor, BMS (**Figure 1D**). Akin to the THP-1 NF-kB reporter results (**Figure 1B**), LCP at 10 μM reduced TNFα-induced IL-8 expression, confirming its anti-inflammatory potential, whereas, at the conditions tested, no impact on IL-8 production was observed upon pre-incubation with the other STLs tested (**Figure 1D**).

The lack of inhibitory effect observed DHLC and DHLCP may be explained by structural differences among the tested compounds. Specifically, while LC and LCP possess the α-methylene-γ-butyrolactone moiety, which is a key pharmacophore conferring electrophilic reactivity and enabling covalent interactions with nucleophilic residues in target proteins ^41^, their hydrogenated derivatives lack this α-methylene group as a result of reduction of the exocyclic double bond. This modification abolishes Michael acceptor reactivity and may contribute to the reduced or absent anti-inflammatory response observed for DHLC and DHLCP under the tested conditions.

The effect on NF-kB and the reduction of IL-8 in endothelial cells upon LCP exposure may reflect a potential impact on systemic inflammation and its effects. In fact, an elegant study by He et al. demonstrated that LCP inhibits the NF-kB-mediated inflammatory response in ApoE–/– mice, by a mechanism involving the prevention of cytoplasmic dynein-mediated translocation of p65 in inflamed macrophages, thereby delaying the development of atherosclerosis^42^. Consequently, and focusing on the impact of LCP on NF-kB in human endothelial cells, we generated an NF-kB luciferase reporter EA.hy926 cell line (**Figure 2A**). As observed in THP-1 macrophages, TNFα exposure induced NF-kB activation, which could be blocked in the presence of LCP (5 and 10 µM), being as effective at 10 µM as the NF-kB inhibitor control, BMS (**Figure 2A**). Importantly, LCP at 10 µM did not affect cell viability (**Supporting information; Figure S2**). Since the higher effects were observed at 10 µM, we used this concentration of LCP in subsequent experiments. To further confirm the impact of LCP on NF-kB in EA.hy926 cells, and following the prophylactic approach (**Figure 1C**), we assessed NF-kB nuclear translocation, as a readout of NF-kB activation^43^.

**Figure 2.**
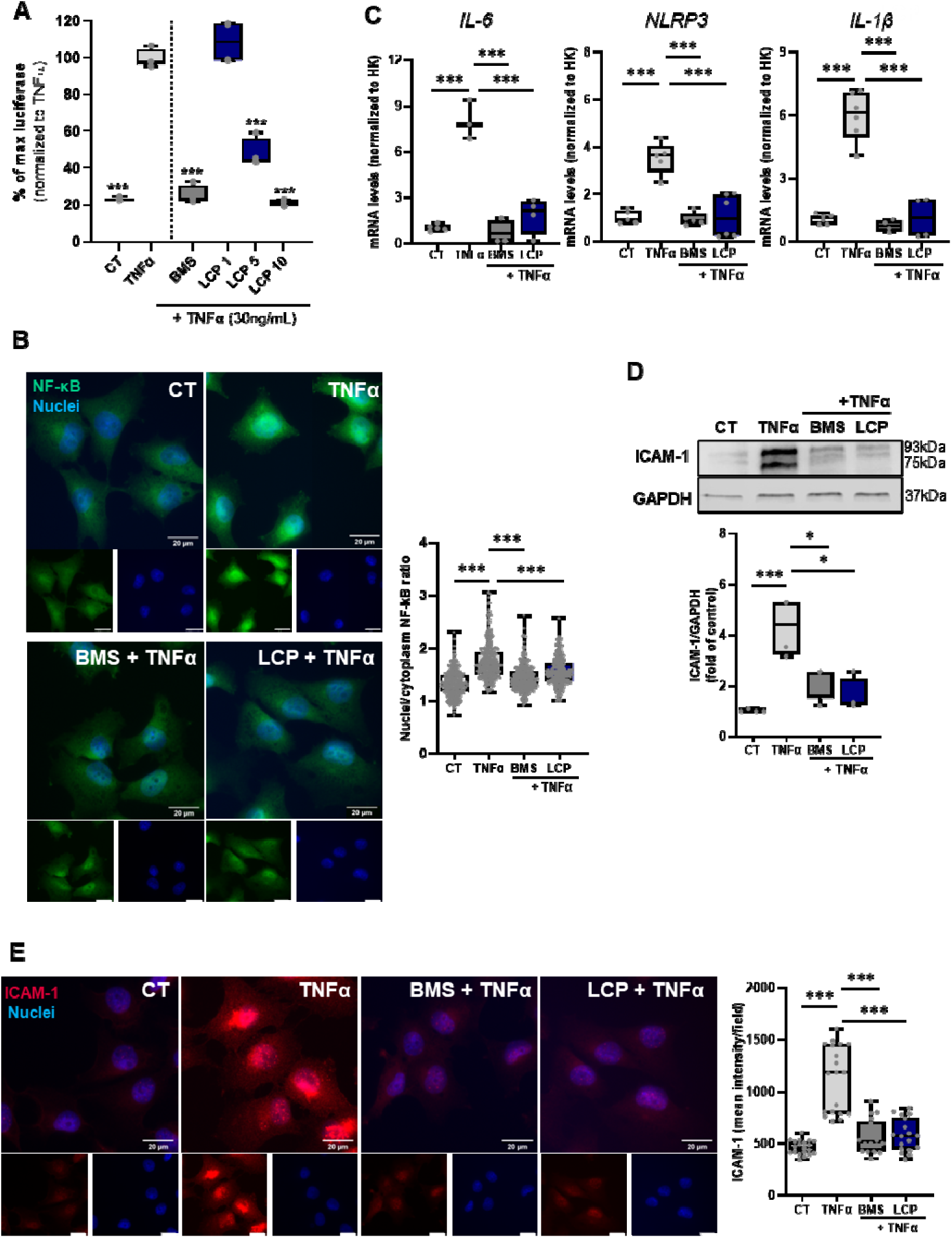
Lactucopicrin impacts the NF-kB signaling pathway in human endothelial cells (EA.hy926). **A)** EA.hy926 NF-kB luciferase reporter cells readout upon 4 h exposure to TNFα (30 ng/mL) in the absence (left) or presence of different concentrations of LCP (1, 5, and 10 μM, right) for 3h. **B**) Immunofluorescence analysis, NF-kB expression labeled in green and nuclei in blue. Left panel: Representative images from three independent biological replicates are shown. Scale bar: 20 μm. Right panel: quantification of NF-kB p65 nuclear translocation. At least 20 images per condition were analyzed, with approximately 15 cells per image (per each replicate). Results are presented as box plots summarizing the distribution of nuclear translocation values obtained from individual cells. EA.hy926 were pre-incubated with exposure to 0.5% DMSO (CT), BMS (5 μM), or LCP (10 μM) for 1 hour, followed by 1h exposure to TNFα (30 ng/mL). Each dot represents values in each analyzed cell. **C**) RT-qPCR gene expression analysis of *IL-6*, *NLRP3,* and *IL-1*β of EA.hy926 cells collected upon 1h pre-treatment with DMSO (CT), BMS (5 μM), or LCP (10 μM), followed by 3 h exposure to TNFα (30 ng/mL). The RT-qPCR data are normalized for the expression of the housekeeping (HK) genes, β-actin and HPRT1, and presented as a relative gene expression normalized to CT (DMSO). **D)** Western blot analysis of ICAM-1 protein, GAPDH used as loading control. EA.hy926 cells were collected upon 1h pre-treatment with DMSO (CT), BMS (5 μM), or LCP (10 μM), followed by 3 h exposure to TNFα (30 ng/mL). **E)** Immunofluorescence analysis, ICAM-1 expression labeled in red and nuclei in blue. Left panel: Representative images from three independent experiments are shown. Scale bar: 20 μm. Right panel: quantification of ICAM-1 expression. At least 4 images per condition with 10 cells per image (for each replicate) were analyzed. Results are presented as box plots summarizing ICAM-1 expression levels per image. Each dot represents the mean of all detected spots per image. Data are shown as the mean ± SD. One-way ANOVA Dunnett post hoc was used to evaluate the significant differences between treatments and TNFα, *** *p*<0.001 or \**p*<0.05.

As depicted in **Figure 2B**, TNFαLinduced translocation of NF-kB to the nucleus, which could be reverted in the presence of the NF-kB inhibitor, BMS. Similarly, LCP impacted TNFα-induced NF-kB nuclear translocation, further confirming its antagonistic effect on TNFα-elicited NF-kB activation.

The NF-kB signaling pathway regulates a vast number of inflammatory mediators, including cytokines, chemokines, and other molecules^44^. We performed gene expression analysis of different NF-kB targets, as described to be involved in endothelial dysfunction, namely *IL-6*, *NLRP3,* and *IL-1*β^45,46^. The expression of the pro-inflammatory cytokines (*IL-1*β and *IL-6*) and *NLRP3* (cytosolic pattern recognition receptor, inflammasome component) was significantly increased upon stimulation of EA.hy926 cells with TNFα (**Figure 2C**). Albeit similar to BMS, co-exposure to LCP significantly reduced their TNFα-induced levels (**Figure 2C**). In addition, we evaluated the expression of ICAM-1 (Intercellular Adhesion Molecule-1), known to play a crucial role in inflammatory contexts within endothelial cells^47^. ICAM-1 acts as a cell surface glycoprotein involved in various processes related to inflammation, including leukocyte recruitment and adhesion to the endothelium^48^, and its upregulation is associated with CVD^49^. Noteworthy, NF-kB mediates the expression of ICAM-1 in response to injury or inflammatory stimuli, such as TNFα. Consistently, our results showed a significant increase in ICAM-1 expression in EA.hy926 cells upon TNFα stimulation (**Figure 2D**). Co-treatment of cells with either BMS or LCP reduced the TNFα-induced ICAM-1 expression (**Figure 2D**). These results were further corroborated by immunofluorescence detection of ICAM-1 (**Figure 2E**). In control cells, ICAM-1 expression is typically low (**Figure 2E**). However, following TNF-α stimulation, a strong increase in fluorescence intensity throughout the cell and over the nuclei was observed, an effect abolished by LCP, as well as by BMS (**Figure 2E**).

Our results demonstrate that LCP effectively mitigates inflammation-induced endothelial dysfunction, as evidenced by the prevention of NF-kB nuclear translocation in these cells and by the reduction in TNF-α-induced ICAM-1 and pro-inflammatory mediator expression in response to inflammatory stimuli. The modulation of ICAM-1 expression by LCP is particularly relevant because it plays a central role in the early stages of endothelial dysfunction by mediating the adhesion and transmigration of leukocytes across the endothelium into tissues during inflammation, a critical event in the initiation and progression of atherosclerosis^47^. Its sustained expression under pro-inflammatory conditions promotes vascular inflammation, increases endothelial permeability, and exacerbates vascular damage. Thus, the ability of LCP to prevent TNFα-induced ICAM-1 upregulation highlights its potential to preserve endothelial integrity and attenuate leukocyte recruitment, positioning LCP as a promising candidate to counteract vascular inflammation and atherosclerotic disease progression.

Importantly, TNFα is a well-known pro-inflammatory cytokine that is commonly found in atherosclerotic lesions^50^. This cytokine provides cell signals leading to the activation of NF-kB, and subsequent activation of downstream genes like *IL-6* and *IL-1*β in cardiovascular pathologies^51^, all of which expression was reduced by LCP exposure in our experiments, akin to what observed in the presence of the chemical inhibitor of NF-kB activation, BMS^40^.

Overall, our results show that LCP can modulate NF-kB activity in both macrophages and endothelial cells. Previous studies have evaluated the anti-inflammatory potential of chicory-derived STLs through other molecular pathways. In particular, 11β,13-dihydrolactucin was shown to modulate inflammatory responses through the modulation of other pathways, specifically the nuclear factor of activated T-cells (NFAT) using a yeast reporter system^52^. Although that study evaluated the cytotoxicity of all the STLs tested in our current work, it did not assess LCP specifically or directly compare the effects of individual STLs. To our knowledge, LCP has not been previously investigated in this context, and our findings provide novel insights into its specific role in regulating inflammation via the NF-kB signaling pathway.

### LCP as an AHR antagonist

The identification of a small molecule capable of targeting both NF-kB and AHR is of significant importance due to the intricate relationship between these signaling pathways and their roles in regulating immune responses and inflammatory processes^53–57^. We took advantage of a previously established *in silico* modeling approach (Induced Fit Docking (IFD) protocol as described elsewhere ^18,19,35,36^ to evaluate the potential of the different STLs as AHR ligands. As shown in **Figures 3A** and **3 B**, the predicted best poses for the tested STLs to fit into the AHR ligand binding domain were obtained for LCP (lower IFD scores). Interestingly, LCP and DHLCP are predicted to form a similar interaction with the backbone carboxyl of I349 and the aromatic ring of F351 (**Figure 3A**). While LC and DHLC do not extend that far due to the lack of the phenolic ring. In general, the positioning of the core structure of the compounds to human AHR was similar. However, the hydrogen bonding varied for each compound, which might account for the different modulatory properties (**Figure 3B**). The best docking pose was obtained for LCP, resulting in a lower IFD score compared to the other STLs.

**Figure 3.**
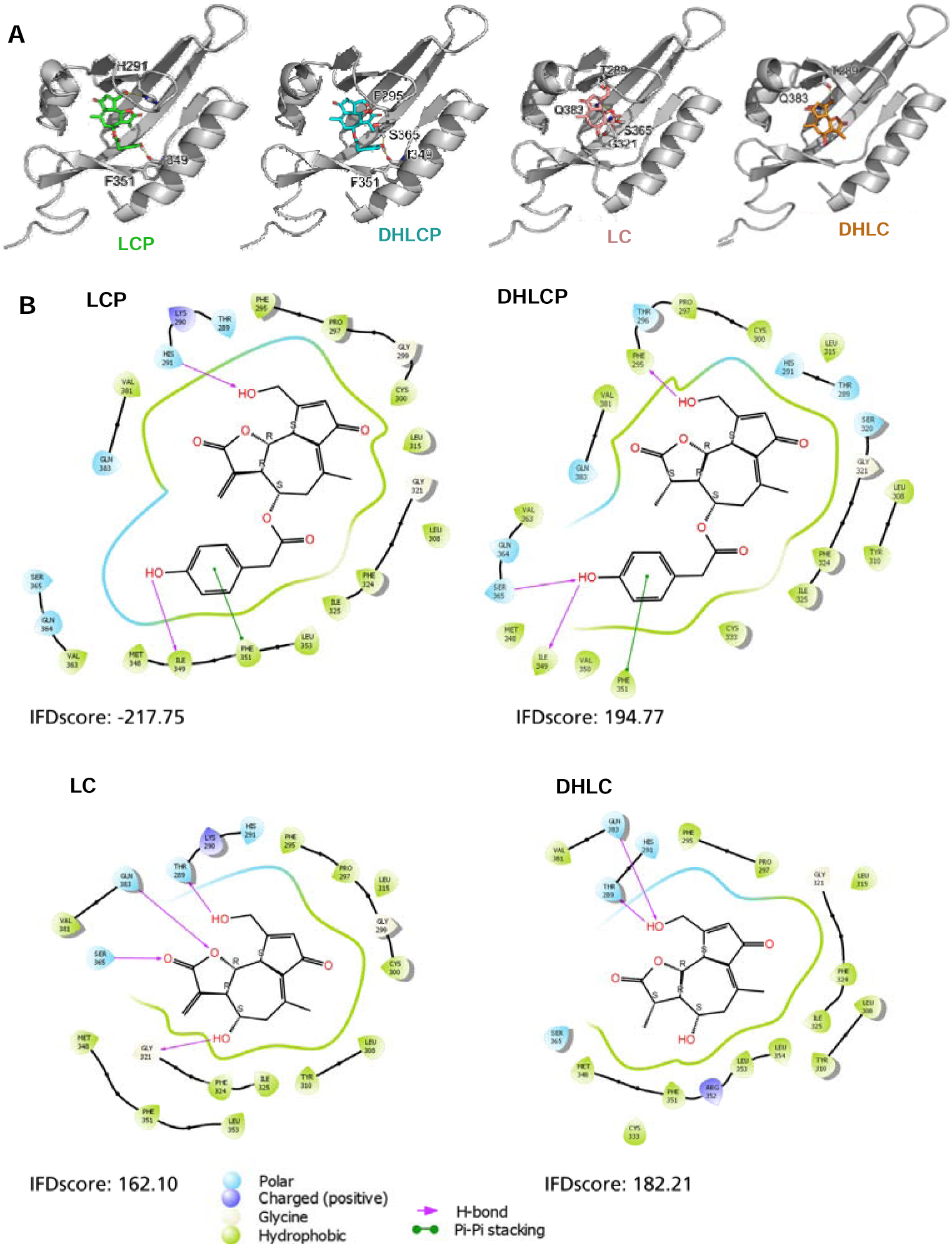
*In silico* modelling for lactucopicrin (LCP), 11β,13-dihydrolactucopicrin (DHLCP), lactucin (LC), and 11β,13-dihydrolactucopicrin (DHLCP) with human AHR. **A)** Docking pose in the human AHR (cartoon, grey). Residues predicted to form hydrogen bonds to the compounds or to be involved in aromatic pi-pi stacking are shown as sticks. **B)** 2D-ligand interaction diagram of the best induced fit docking poses of the 4 STLs in the human AHR. Hydrogen bonds are depicted as pink arrows, and pi-pi stacking is depicted as green lines. Arrows originating from or ending at the pointy side of amino acids indicate side-chain interactions, and the blunt side depicts the amino acid backbone.

To confirm whether the different STLs and LCP in particular modulate the AHR signaling pathway, we resorted to a well-established THP-1 AHR luciferase reporter system^18,19,35–37^. AHR activation in THP-1 observed upon exposure to indigo, a known AHR agonist^16,20^ (**Figure 4A, left**) is not mimicked by exposure to the STLs (**Supporting information; Figure S3**). Albeit, similar to what was performed for the NF-kB, we evaluated whether the STLs could impact AHR activation induced by indigo^17,20^. Exposure to LCP, and to a lesser extent to LC (observed only at the highest concentration tested, 100 µM), was able to antagonize the AHR activation elicited upon exposure to indigo (1 µM; **Figure 4A**).

**Figure 4.**
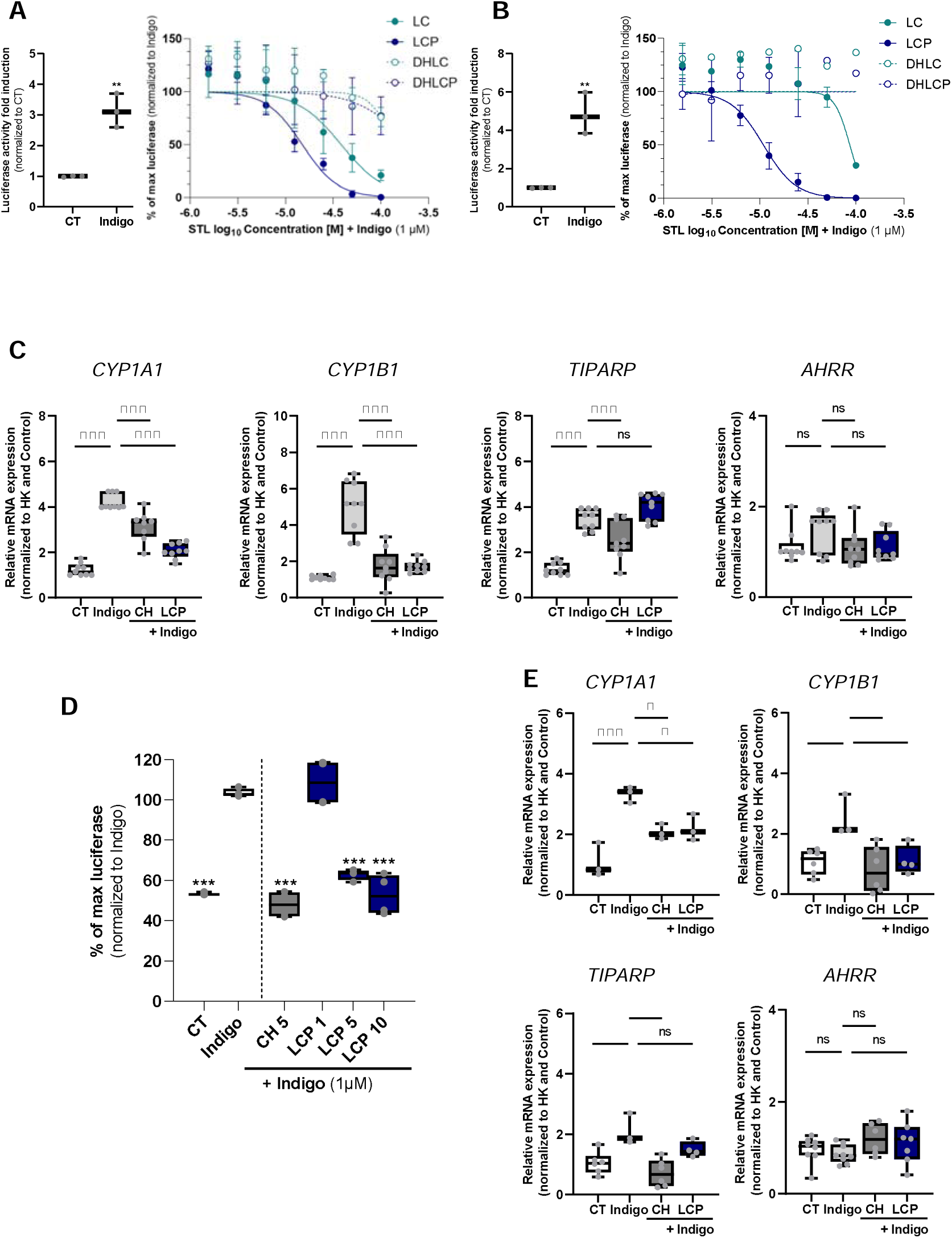
Lactucopicrin (LCP) as an AHR antagonist. **A)** THP-1 (human monocytic) AHR luciferase reporter cells readout upon 4h exposure to indigo (1 µM, left) or to different concentrations of STLs (right), lactucin (LC), lactucopicrin (LCP), dihydrolactucin (DHLC), or dihydrolactucopicrin (DHLCP) in the presence of indigo (1 µM). **B)** Caco-2 (human colon adenocarcinoma) AHR luciferase reporter cells readout upon 4h exposure to indigo (1 µM, left) or to different concentrations of STLs (right) in the presence of indigo (1 µM). **C)** RT-qPCR gene expression analysis of *CYP1A1*, *CYP1B1*, *TIPARP,* and *AHRR* on Caco-2 cells collected upon 1h pre-treatment with DMSO (CT), BMS (5 μM), or LCP (12.5 μM) followed by 3 h exposure to indigo (1µM). The RT-qPCR data are normalized for the expression of the housekeeping (HK) genes, β*-actin* and *HPRT1*, and presented as a relative gene expression normalized to CT (DMSO). **D**) EA.hy926 AHR luciferase reporter cells readout upon exposure to LCP (1, 5, and 10 μM) or CH-223191 (CH, 5 µM) for 1h, followed by 3h exposure to indigo (1 µM). **E**) RT-qPCR gene expression analysis of *CYP1A1*, *CYP1B1*, *TIPARP,* and *AHRR* on EA.hy926 cells collected upon 1h pre-treatment with DMSO (CT), BMS (5 μM), or LCP (12.5 μM), followed by 3 h exposure to indigo (1µM). The RT-qPCR data are normalized for the expression of the HK genes, β*-actin* and *HPRT1*, and presented as a relative gene expression normalized to CT (DMSO). Data are shown as the mean ± SD. One-way ANOVA with Dunnett post hoc test was used to evaluate the significant differences between treatments and Indigo, \*\*\**p* < 0.001, \*\**p* < 0.01, \**p* < 0.05, or ns (non-significant).

Since modulation of the AHR can vary depending on the ligand, cellular, and environmental context, ^17,20,54,58^, we decided to explore the impact of these STLs on the AHR using other human cell lines. AHR plays a crucial role in the intestine, contributing to the detoxification of xenobiotics, immune regulation, maintenance of barrier integrity, interactions with the gut microbiota, and influencing stem cell function, making it particularly relevant to intestinal health^59^. Thus, we decided to assess their impact on a human colon epithelial cell line (Caco-2, a human colon adenocarcinoma cell line), establishing an AHR luciferase reporter in this cell line as well. Similar to THP-1 cells, LCP and, to a lesser extent, LC antagonized the AHR activation elicited by exposure to indigo in Caco-2 cells, whereas, under the tested conditions, DHLC and DHLCP did not (**Figure 4B**).

To gain more insights into the effects of LCP as an AHR antagonist, we measured by RT-qPCR the expression of different AHR target genes, namely the detoxifying monooxygenases (*CYP1A1* and *CYP1B1*)^60^, the AHR repressor (*AHRR*), and the downstream effector TCDD-inducible poly(ADP-ribose) polymerase (*TIPARP*)^61^. Under the tested conditions, indigo (1 µM) induced the expression of *CYP1A1*, *CYP1B1,* and *TIPARP* in Caco-2 cells, whereas no statistically significant differences were observed for *AHRR* (**Figure 4C**). Notably, we and others have demonstrated that induction of different AHR target genes, including AHRR, does not always correlate with the expression of *CYP1A1* and *CYP1B1* and that differences between ligands, cell type, and exposure conditions occur^62^. Remarkably, LCP was able to block the induction of *CYP1A1* and *CYP1B1,* but not *TIPARP* gene expression, a pattern shared with the positive control, the AHR inhibitor CH223191 (CH)^63^ (**Figure 4C**).

Considering the results obtained with LCP in EA.hy926 cells, its effects on NF-kB, and its potential use as an anti-inflammatory agent in this context, we also evaluated its impact on AHR activity in these cells. Therein, and to the same levels of the AHR inhibitor CH, LCP was able to inhibit indigo-induced AHR activity, as measured using an AHR luciferase assay (**Figure 4D**) and reduce the gene expression of the AHR target genes *CYP1A1* and *CYP1B1* (**Figure 4E**). To further validate the impact of LCP on AHR modulation, we assessed CYP1A1 activity using an ethoxyresorufin-O-deethylase (EROD) enzymatic assay^18,19^ on Caco-2 cells. As occurred for CH, LCP exposure blocked CYP1A1 enzymatic activity induced by indigo (**Supporting information; Figure S4**)

In light of our findings, which show the ability of LCP to modulate both NF-kB and AHR pathways simultaneously, underscores its unique therapeutic potential, as it targets two pivotal regulators of immune and inflammatory responses, offering a promising dual-action strategy for controlling inflammation in complex disease contexts.

Beyond its classical role in xenobiotic sensing, the AHR signaling axis has been increasingly recognized as a key modulator of host-pathogen interactions, mucosal immunity, and inflammatory homeostasis ^54^. In viral infections, AHR activity modulates the host response by balancing immune activation and tissue protection^64^. Similarly, in the context of bacterial and parasitic infections, AHR has been shown to fine-tune inflammatory signaling, promoting pathogen clearance while preventing excessive tissue damage^18,19^. In this line, we have previously demonstrated that LCP exhibits antiviral activity both in silico and in vitro against two key proteases of SARS-CoV-2^65^.

However, despite growing evidence highlighting AHR as a promising immunomodulatory target, the therapeutic potential of natural small molecules to manage inflammation via AHR modulation has remained largely unexplored until now. Except for tapinarof, which is a natural AHR agonist that resolves skin inflammation in mice and human^23,66–68^ evidence for dual-target therapies by small molecules affecting both pathways is scarce. Some flavonoids, curcumin, and resveratrol are reported to interact with AHR and also exhibit anti-inflammatory activities; however, the interconnection between both pathways mediated by the compounds has not been fully established^20,69^. Therefore, LCP emerges as a novel small molecule with this dual role, offering a strategic advantage in targeting interconnected immune and inflammatory mechanisms for more effective disease control.

### Anti-inflammatory effects of LCP targeting the crosstalk between AHR and NF-kB signaling pathways

Several studies have shown that AHR antagonists can attenuate inflammation in various cell types.^70,71^ Previous evidence has described that chemical AHR inhibitors block the NF-kB signaling pathway, suggesting this crosstalk as the underlying mechanism for the decrease in the peripheral and central inflammatory processes^72,73^. Therefore, to validate the LCP dual role, dissect its effect on AHR and NF-kB crosstalk, and to further investigate the potential mechanism responsible for the anti-inflammatory effect of LCP, we established an AHR knockdown (AHR-KD) EA.hy926 cell line (**Figure 5A**). Following the same prophylactic approach as previously, we measured IL-8 production in AHR-KD endothelial cells after incubation with LCP and BMS. As previously observed, BMS or LCP exposure blocked TNFα-induced IL-8 expression in wild-type (AHR-WT) EA.hy926 cells (**Figure 1D**). However, whereas this effect was still observed in BMS-treated AHR-KD cells, the LCP blocking effect was lost (**Figure 5B**). To further analyze this effect, we evaluated the expression of *IL-6*, *NLRP3,* and *IL-1*β in both AHR-WT and AHR-KD cells under identical treatment conditions. As expected, TNFα increased the levels of these genes in both AHR-WT and AHR-KD cells. Strikingly, whereas BMS reduced the TNFα-induced levels of all the genes assayed in both AHR-WT and AHR-KD cells, the LCP impact on reducing the expression of these genes in WT cells was lost in AHR-KD cells (**Figure 5C)**, confirming a pivotal role for AHR in mediating the LCP-elicited effects.

**Figure 5.**
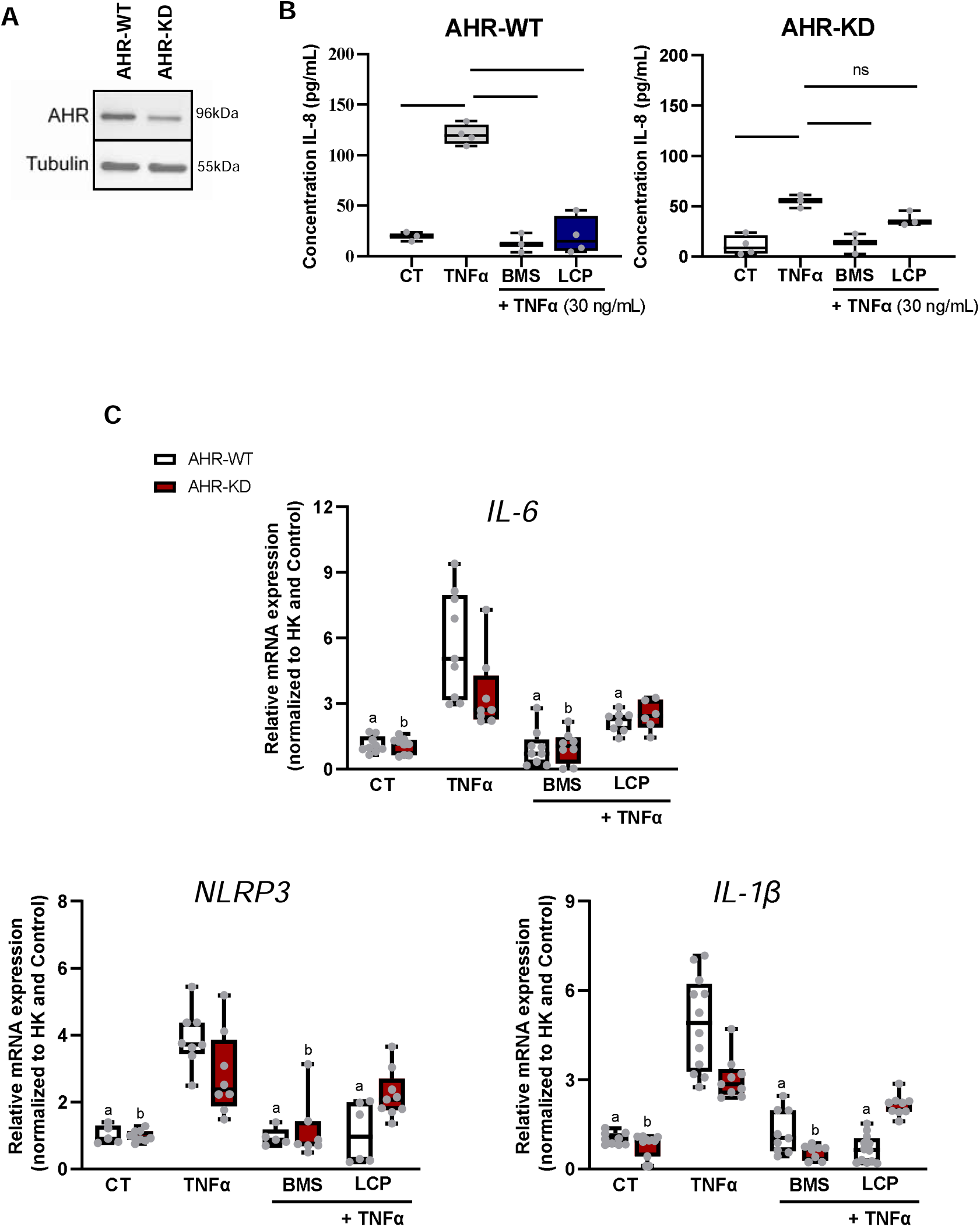
LCP modulates the crosstalk between AHR and NF-kB. **A)** AHR protein expression detection by Western blot in wild type (AHR-WT) and AHR knockdown (AHR-KD) EA.hy926 cells. Tubulin was used as a loading control. **B**) EA.hy926 endothelial cells (AHR-WT or AHR-KD) IL-8 release upon exposure to TNFα or TNFα in the presence of LCP (10 µM) or the NF-kB inhibitor BMS-345541 (BMS, 5 µM). **C)** RT-qPCR gene expression analysis of *IL-6*, *NLRP3,* and *IL-1*β of EA.hy926 cells (AHR-WT or AHR-KD) collected upon 1h pre-treatment with DMSO (CT), BMS (5 μM), or LCP (10 μM), followed by 3 h exposure to TNFα (30 ng/mL). Data are shown as the mean ± SD. One-way ANOVA with the Dunnett post hoc test was used to evaluate the significant differences between treatments and TNFα, \*\*\**p* < 0.001 or ns, non-significant. ^a^ Represents significant differences between TNFα+ treatments vs TNFα in AHR-WT cells. ^b^ Represents the significant differences between TNFα+ treatments vs TNFα-in AHR-KD cells.

It is known that AHR regulates the expression of inflammatory mediators, including those mediated by its interaction with other key signaling pathways involved in this process, such as the NF-kB pathway.^17,54,56^

Previous reports have established the direct interaction of AHR and NF-kB^21,53,74^, but the exact mechanism(s) of action are not fully understood. Physical interaction between the AHR and NF-kB has been reported, for example, between the AHR and RelB, regulating the expression of different cytokines, such as IL-8^74^. Furthermore, AHR overexpression increases NF-kB activity and enhances the association of the AHR/RelA complex with NF-kB binding site on pro-inflammatory cytokine promoters (*e.g*., IL-6)^75^. Likewise, the interaction between AHR and NF-kB has also been shown to modulate the activity of the AHR^21,74^. RelA is directly involved in the nuclear translocation of NF-kB and, therefore, in the activation of this signaling pathway^76^. Of note, RelA has been shown to mediate LPS-induced AHR expression^21,74^. TNFα induction of AHR expression has also been shown to occur, as well as a cross-regulatory effect on both the expression of AHR target- and NF-kB target-genes (e.g. IL-8, IL-1β, IL-6)^77,78^ in the presence of AHR (e.g. TCDD) and NF-kB modulators (e.g. TNFα). Our results indicate that LCP interferes with this crosstalk, further substantiating this interaction and its involvement in regulating inflammation, as evidenced by the LCP-mediated inhibition of NF-kB signaling after stimulation with TNFα being blocked in cells with reduced AHR expression (AHR-KD; **Figure 5**).

Previous studies have demonstrated that LCP inhibits NF-kB activation in both endothelial cells and macrophages. In endothelial cells, LCP reduced vascular inflammation and improved the response in a mouse model of sepsis^79^. In contrast, in macrophages, LCP impacts NF-kB/p65 nuclear translocation by inhibiting dynein-mediated transport. This mechanism effectively reduces inflammation in atherosclerotic plaques in an ApoE−/− mouse model of atherosclerosis^80^.

Our data reveals LCP unique ability to modulate both the aryl hydrocarbon receptor (AHR) and the nuclear factor kappa B (NF-kB) signaling pathways—two central regulators of immune and inflammatory responses.

## CONCLUSIONS

Our results confirm the impact of the chicory-derived LCP on the regulation of inflammation and further unveil its capacity to modulate the AHR pathway and the crosstalk between the AHR and NF-kB. Over the last decade, numerous endogenous and natural AHR modulators have been identified, including microbiota-derived molecules, tryptophan, and kynurenic acid metabolites, as well as plant-derived compounds, many of which serve as dietary AHR agonists or antagonists. Due to different AHR functions and its regulated responses, which can vary according to the ligand and cellular and tissue contexts, the identification of new natural AHR modulators is still of major interest, as new ligands can aid in modulating a specific AHR function and be further explored as part of host-directed therapeutic approaches targeting the AHR. The herein-revealed mechanism of LCP anti-inflammatory effects via the AHR confirms the potential of targeting this receptor as a promising therapeutic strategy for modulating NF-kB activity to mitigate inflammatory responses. Given the crucial roles of NF-kB and AHR in CVD, LCP’s dual-action mechanism positions it as a potential candidate for the treatment and prevention of vascular inflammation and related cardiovascular conditions. This dual modulatory capacity positions LCP as a compelling candidate for the development of next-generation anti-inflammatory therapies. By simultaneously targeting AHR, which governs immune homeostasis and tolerance, and NF-kB, a master regulator of pro-inflammatory gene expression, LCP offers a synergistic mechanism to dampen excessive inflammation while preserving immune balance. Such a dual-action strategy is especially valuable in complex inflammatory diseases, where single-pathway interventions often fall short. The ability of LCP to engage both pathways not only enhances its therapeutic potential but also supports the broader concept of multi-targeted approaches in inflammation control, particularly when derived from safe, bioactive natural sources. Given that LCP is a molecule of dietary relevance, its translational potential is exceptionally high. Therefore, rigorous in vivo validation of its immunomodulatory effects is not only warranted but essential to fully harness its promise as a safe and effective anti-inflammatory therapeutics crucial for several inflammatory diseases, including cardiovascular dysfunctions. Although this work provides proof-of-concept mechanistic evidence in human cell-based systems, further validation in more complex models, organoid or in vivo models, will be required to confirm its translational relevance. Any future in vivo assessment should be preceded by toxicity and pharmacokinetic studies to establish safe and effective dosing. These steps will be essential to substantiate the therapeutic potential of LCP as a multi-target anti-inflammatory agent.

## EXPERIMENTAL SECTION

### Materials

Lactucin (LC), lactucopicrin (LCP), 11β-13-dihydrolactucin (DHLC) and 11β-13-dihydrolactucopicrin (DHLCP) were acquired from Extrasynthese (3809, 3813, 3810, 3811) (Genay Cedex, France). Indigo (CAS No. 482-89-3) was obtained from BOC Sciences (Shirley, NY, USA). CH-223191 (CH) was purchased from Sigma-Aldrich (St. Louis, MO, USA). BMS-345541 (BMS, CAS No. 547757-23-3) was acquired from Selleck Chemicals (Planegg, Germany). Tumor necrosis factor-alpha (TNFα) was purchased from PeproTech (London, United Kingdom). Milli-Q water purification system (Merck Millipore, Billerica, MA, USA) was used in all experiments.

### Cells

EA.hy926 human endothelial cells, THP-1 human macrophage cells, and Caco-2 human colon cancer cells were obtained from the American Type Culture Collection (ATCC, Manassas, VA, USA). EA.hy926 and Caco-2 cells were cultured in Dulbecco’s Modified Eagle’s Medium (DMEM; GIBCO, 31966047) supplemented with 10% (v/v) Fetal Bovine Serum (FBS; GIBCO, 16140071), and THP-1 cells cultured in RPMI 1640 (GIBCO, 31870074) supplemented with 10% (v/v) FBS (GIBCO, 16140071), 1% (v/v) L-Glutamine (GIBCO, 25030081) 1% (v/v) non-essential amino acids (Cytiva, 15333581), 1% (v/v) Sodium pyruvate (Cytiva, 11501871), 1% (v/v) HEPES (Cytiva, 10204932) and 0,05 mM 2-mercaptoethanol (GIBCO, 31350010).

### Lentiviral production

Lentiviruses were produced according to the described TRC lentiviral proceedings (https://portals.broadinstitute.org/gpp/public/resources/protocols). Briefly, HEK293T packaging cells were seeded at a density of 2×10^5^ cells/mL in DMEM high glucose complete medium (Cytiva) in 96-well plates. After 24h incubation, cells were transfected with a lentiviral packaging mix (Sigma-Aldrich) and 100 ng of the respective CRISPR lentiviral vector containing pLV-U6g-EPCG vector (Sigma Aldrich, AHR gRNA targeting sequence - AGTCGGTCTCTATGCCGCTTGG), using Fugene 6 (Roche, Berlin, Germany) in Optimem medium (Gibco). After 18h of incubation, the medium was carefully aspirated and replaced with a high serum growth cell culture medium (DMEM+ 30% FCS (v/v)). Viruses were harvested at 48h and 72h post-transfection. Viral supernatants were centrifuged at 2100 rcf for 5 minutes, filtered through a 0.45 µm PES filter, and stored at -80°C until further use.

The lentiviral constructs (CLS-2045L and CLS-013L) for generating the AHR and NF-kB reporter cell lines, respectively, were obtained from QIAGEN.

### Lentiviral transduction to generate luciferase reporter and knockdown cells

Lentiviral transduction was performed as described previously^18,19,35,36^ and according to the protocols available at https://portals.broadinstitute.org/gpp/public/resources/protocols. In brief, cells were seeded at a density of 2.2x104 cells per well (for Caco-2 or EA.hy926 the day before infection) or 5x10^4^ cells per well (for THP-1 the day of infection) in a 96-well plate. Lentiviruses were added to the cells in a medium containing 8 µg/ml polybrene (Sigma-Aldrich). Plates were spinoculated for 90 min at 2200 rpm at 37°C. At 2 days post-infection, transduced cells were selected using Puromycin (Calbiochem; 5 mg/ml). For CRISPR-KD cell line generation, based on GFP expression (CRISPR constructs carry GFP for selection), cells were single-cell sorted (FACSAria II, BD Biosciences) into 96-well plates and further expanded.

### Luciferase activity measurements

Luciferase activity measurements were performed as a readout for AHR or NF-kB activation, using established luciferase reporter cells (THP-1 AHR-, THP-1 NFkB-, or Caco-2 AHR-luciferase reporters) or herein generated (EA.hy926 NF-kB luciferase reporter, see details above^18,19,36,37,81^. Of note, FICZ was used as an agonist control for the AHR reporter cells, whereas TNFα was used as an NF-kB agonist control^18,19,36^. THP-1 reporter cells were differentiated into macrophages following a well-established protocol^18,81^. In brief, 5x10^4^ cells per well were plated in a 96-well plate and treated with 200 nM phorbol 12-myristate 13-acetate (PMA) for three days. This was followed by washing with PBS and then incubating the cells for four days in complete RPMI medium (Cytiva) before challenging them with different ligands. EA.hy926 and Caco-2 reporter cells were plated at a density of 2x10^4^ cells per well in a 96-well plate in their respective growing medium for 24h, before the addition of the different ligands. After the specified conditions (ligand type, ligand concentration, and exposure time), the cells were rinsed with PBS and harvested in a lysis reagent (Cat. # E1531, Promega). The lysates were used to determine luciferase activity using the Luciferase Assay System (Cat.#E1501, Promega) according to the manufacturer’s instructions. Luciferase activity was normalized to the total concentration of protein in solution determined by bicinchoninic acid assay (Pierce BCA Protein Assay, Cat. #23227, Thermo Scientific). Results are shown as fold induction determined upon normalization to the luciferase values of the respective control.

### Cytokine analysis

EAhy926 endothelial cells were cultured in Dulbecco’s modified Eagle’s medium with 10 % fetal calf serum. For treatment experiments, cells were seeded for 48 h. Briefly, cells were seeded in 24-well plates at a density of 8x10^4^ cells/well. After 48h, cells were pre-treated with different STLs (1, 5, or 10 µM) or BMS (5 µM) for 1 h, before an additional 3h in the presence of an inflammatory stimulus with TNFα (30 ng/mL). Cell supernatants were collected and stored at −80 °C until further analysis. IL-8 release induced by TNFα in EA.hy926 endothelial cells was assessed by enzyme-linked immunosorbent assay (ELISA) according to the manufacturer’s instructions (KHC0081; Invitrogen, Camarillo, CA, USA). The plates were incubated at room temperature in the dark for 30 minutes, using a Synergy HT microplate reader (Biotek®, Winooski, USA), and the absorbance was measured at 450nm. All the plates and reagents were included in the kit.

#### EROD assay

The EROD assay was used to detect the CYP1A1 enzymatic activity by measuring the conversion of ethoxyresorufin to resorufin^18,19,35,36,82^ in Caco-2 cells (4x10^4^ cells/well in 96-well plates) exposed for 24h to different ligands. Briefly after stimulation of the cells with the diverse ligands, 4 µM resorufin ethyl ether (EROD, Sigma-Aldrich) and 10 µM dicoumarol (Sigma-Aldrich) were added to the cell culture for 1h, followed by measuring the resorufin fluorescence using a Spectramax ID3 (Molecular Devices) or a Tecan Sparks (Tecan) microplate reader. The activity was corrected to the amount of protein, measured by Pierce BCA Protein Assay (Cat. #23227, Thermo Scientific), and normalized to the respective control.

### Reverse Transcription–qPCR (qPCR)

For NF-kB treatments, after 48h, cells were pre-treated with lactucopicrin (10 µM) or BMS (5 µM) for 1 h before exposure to the inflammatory stimulus (TNFα, 30 ng/mL) for 3 h. For AHR treatments, cells were pre-treated with lactucopicrin (10 µM) and CH-223191 (5 µM) for 1 h before the induction of AHR with Indigo (1 µM) for an additional 3 h. The qPCR analyses were performed according to MIQE guidelines (Minimum Information for Publication of Quantitative Real-Time PCR Experiments)^83^. Total RNA was extracted using the RNeasy Mini kit (QIAGEN). After cleaning, 130-300 ng of total RNA was used for reverse-transcription with SuperScript™ II Reverse Transcriptase (Invitrogen). The qPCR was performed in a QuantStudio™ 5 (Applied Biosystems), using SensiFAST™ SYBR Lo-ROX Kit (Bioline) to evaluate expression of AHRR (5’-CAAATCCTTCCAAGCGGCATA-3’; 5’-CGCTGAGCCTAAGAACTGAAAG-3’); CYP1A1 (5’-ACATGCTGACCCTGGGAAAG-3’; 5’-GGTGTGGAGCCAATTCGGAT-3’); CYP1B1 (5’-GGGACCGTCTGCCTTGTATG -3’; 5’-GGTGGCATGAGGAATAGTGACA-3’); TIPARP (5’-AATTTGACCAACTACGAAGGCTG-3’; 5’-CAGACTCGGGATACTCTCTCC-3’); IL-6 (5’-ACTCACCTCTTCAGAACGAATTG-3’; 5’-CCATCTTTGGAAGGTTCAGGTTG-3’); IL-1β (5’-AAACAGATGAAGTGCTCCTTCCAGG-3’; 5’-TGGAGAACACCACTTGTTGCTCCA-3’) and NLRP3 (5’-CACCTGTTGTGCAATCTGAAG-3’; 5’-GCAAGATCCTGACAACATGC-3’) genes.

As reference genes were used β-Actin (5’-AACTACCTTCAACTCCATCA-3’; 5’-GAGCAATGATCTTGATCTTCA-3’) and HPRT1 (5’-CCTGGCGTCGTGATTAGTGA-3’; 5’-CGAGCAAGACGTTCAGTCCT-3’). Standard curves were constructed for each gene, and the AbiQuantStudio program was used to extract the quantification cycle (Cq) data. Gene expression was then calculated using the relative quantification method with efficiency correction, as described by the Pfaffl method^84^. The results were expressed as fold-change mRNA levels relative to the control (mRNA fold change) of at least three independent biological replicates.

### Western Blot Analysis

EA.hy926 were seeded in 24-well plates at a density of 8x10^4^ cells/well. After 48h, cells were pre- treated with lactucopicrin (10 µM) or BMS (5 µM) for 1 h before the inflammatory stimulus with TNFα (30 ng/mL), for 3 h. The cells were harvested by briefly washing them with cold PBS, followed by the addition of Cell Lysis Buffer (Cell Signaling Technology). Insoluble material was removed by centrifuging and discarding the pellet. Protein concentration was determined by a Micro BCA™ Protein Assay (ThermoFisher Scientific™). Cell protein extracts (20 µg) were denatured in sample buffer containing 5% 2-β-mercaptoethanol, to be further subjected to a 12% acrylamide SDS-PAGE, transferred to 0.2 µm pore nitrocellulose membranes (#1704159, Bio-Rad) in a Trans-Blot Turbo Transfer System (Bio-Rad), blocked for 1 h with 5% BSA in Tris-Tween buffered saline (TTBS), and incubated overnight under gentle shaking, at 4 °C, with CD54/ICAM-1 (E3Q9N) XP^®^ Rabbit mAb (#67836, Cell Signaling Technology) (diluted 1:1000). Membranes were then incubated at room temperature for 1 h in peroxidase-conjugated secondary antibody: Anti-Rabbit IgG produced in goat (A6154, Sigma-Aldrich) (diluted 1:5000). Membranes were developed using Amersham™ ECL™ Prime Western Blotting Detection Reagent (Cytiva) and visualized using an Odyssey^®^ XF Imaging System (LI-COR^®^). Subsequently, the membranes were incubated with GAPDH Loading Control mAb (MA5-15738-D800, ThermoFisher Scientific™) (diluted 1:5000) and revealed using the same LI-COR^®^ system.

The AHR KD validation was performed using AHR primary antibody (sc-133088, Santa Cruz) and Tubulin Loading Control (T6199, Sigma-Aldrich).

### Immunostaining and image analysis

The seeding of EA.hy926 cells was performed using 8x10^4^ cells/well, which were added to coverslips previously placed in 24-well plates. After 48 h of seeding, the cells were treated with the same treatments detailed in the Western Blot section. Since NF-kB nuclear translocation is a process that occurs very rapidly, pre-incubation with the compounds was applied for 1 h, followed by TNFα stimulation for only 1 h, rather than 3 h. Briefly, the medium was discarded, and cells were fixed with 4% paraformaldehyde (PFA) in PBS for 20 minutes at RT. PFA was removed, and cells were washed three times with PBS. The next step was permeabilization of the cells with 0.3% Triton X-100 in PBS for 15 min. Then, before adding the antibodies, the blocking of the cells was performed with 3% bovine serum albumin (BSA) in PBS for 1h at RT. Lastly, the coverslips were incubated overnight at 4 °C in a humidified chamber with a mixture of primary antibodies for NF-kB and ICAM-1 diluted in PBS with 3% BSA. Primary antibodies were Rabbit Polyclonal Anti-NF-kB p65 (C-20) (Santa Cruz Biotechnology, #SC-372) (diluted 1:200) and Mouse ICAM-1 Monoclonal Antibody (1A29) (Invitrogen, #MA5407) (diluted 1:100). After the overnight incubation, the incubation was done with the secondary antibodies for 1 hour at RT in 3% BSA. A mixture of secondary antibodies was Alexa Fluor 488 Goat anti-Rabbit IgG (H+L) (Invitrogen, #A11034) (diluted 1:100) and Alexa Fluor 568 goat anti-mouse IgG (H+L) (Invitrogen, #A11004) (diluted 1:50). Finally, nuclei were counterstained using Invitrogen™ ProLong Gold Antifade Mountant with DAPI. The images were acquired using a Microscope: Zeiss Axio Imager Z2 (Zeiss, Germany). Four fields per condition were acquired and evaluated for semi-quantitative analysis. Immunofluorescence images obtained by fluorescence microscopy were examined using Icy (Institute Pasteur and France BioImaging, Paris, France) and ImageJ (National Institutes of Health, Bethesda, MD, USA) software.

### Statistical analysis

All statistical analysis was performed using GraphPad Prism 9.0 (GraphPad Software Inc., San Diego, CA, USA). Parametrical data were submitted to one-way ANOVA followed by Tukey post-hoc test, and non-parametrical data were under the Kruskal-Wallis test with post-Dunnett’s multiple comparisons, with a significance level fixed at 95% (*p*<0.05). All the results were expressed as the mean ± standard deviation (SD) from at least 3 independent biological replicates.

### Supporting Information

Supporting Information available: Figures S1–S4 (DOCX); Molecular Formula Strings (CSV).

## AUTHOR INFORMATION

### Corresponding Authors

**Pedro Moura-Alves -** i3S*-*Instituto de Investigaclão e Inovaclão em Saúde*, Universidade do Porto, 4200-135 Porto, Portugal; IBMC-Instituto de Biologia Molecular e Celular, Universidade do Porto, 4200-135 Porto, Portugal; Ludwig Institute for Cancer Research, Nuffield Department of Clinical Medicine, University of Oxford, Oxford OX3 7DQ, United Kingdom*.

**Cláudia Nunes dos Santos -** Instituto de Biologia Experimental e Tecnológica (iBET), Av. República, Qta. Marquês, 2780-157 Oeiras, Portugal; iNOVA4Health, NOVA Medical School|Faculdade de Ciências Médicas, NMS|FCM, Universidade Nova de Lisboa, Lisboa, Portugal; NOVA Institute for Medical Systems Biology, NIMSB, Universidade Nova de Lisboa, 1099-085 Lisboa, Portugal

### Authors

**María Ángeles Ávila-Gálvez –** Instituto de Biologia Experimental e Tecnológica (iBET), Av. República, Qta. Marquês, 2780-157 Oeiras, Portugal; iNOVA4Health, NOVA Medical School|Faculdade de Ciências Médicas, NMS|FCM, Universidade Nova de Lisboa, Lisboa, Portugal.

**Catarina J G Pinto –** iNOVA4Health, NOVA Medical School|Faculdade de Ciências Médicas, NMS|FCM, Universidade Nova de Lisboa, Lisboa, Portugal; i3S*-*Instituto de Investigaclão e Inovaclão em Saúde*, Universidade do Porto, 4200-135 Porto, Portugal; IBMC-Instituto de Biologia Molecular e Celular, Universidade do Porto, 4200-135 Porto, Portugal*.

**Carlos Rafael-Pita –** Instituto de Biologia Experimental e Tecnológica (iBET), Av. República, Qta. Marquês, 2780-157 Oeiras, Portugal; iNOVA4Health, NOVA Medical School|Faculdade de Ciências Médicas, NMS|FCM, Universidade Nova de Lisboa, Lisboa, Portugal.

**Inês Pires Silva –** Instituto de Biologia Experimental e Tecnológica (iBET), Av. República, Qta. Marquês, 2780-157 Oeiras, Portugal; iNOVA4Health, NOVA Medical School|Faculdade de Ciências Médicas, NMS|FCM, Universidade Nova de Lisboa, Lisboa, Portugal.

**Aleksandra T. Janowska –** Ludwig Institute for Cancer Research, Nuffield Department of Clinical Medicine, University of Oxford, Oxford OX3 7DQ, United Kingdom; Cancer Research UK Cambridge Institute, Li Ka Shing Centre, University of Cambridge, Robinson Way, Cambridge, CB2 0RE, UK (present address).

**Sérgio Marinho –** i3S*-*Instituto de Investigaclão e Inovaclão em Saúde*, Universidade do Porto, 4200-135 Porto, Portugal; IBMC-Instituto de Biologia Molecular e Celular, Universidade do Porto, 4200-135 Porto, Portugal*.

**Rory Saitch –** *Ludwig Institute for Cancer Research, Nuffield Department of Clinical Medicine, University of Oxford, Oxford OX3 7DQ, United Kingdom*.

**Yilong Lian –** *Ludwig Institute for Cancer Research, Nuffield Department of Clinical Medicine, University of Oxford, Oxford OX3 7DQ, United Kingdom*.

**Pakavarin Louphrasitthiphol –** *Ludwig Institute for Cancer Research, Nuffield Department of Clinical Medicine, University of Oxford, Oxford OX3 7DQ, United Kingdom*.

**Jonas Protze –** *FMP, Leibniz-Forschungsinstitut für Molekulare Pharmakologie, 13125 Berlin, Germany*.

**Gerd Krause -** *FMP, Leibniz-Forschungsinstitut für Molekulare Pharmakologie, 13125 Berlin, Germany*.

### Author Contributions

JP and GK performed the in silico docking analysis. YL, RS, ATJ, SM, and PL conducted the luciferase reporter assays, EROD activity measurements, and cell viability experiments. YL, PL, and SM generated various cell lines, including luminescence reporter and AHR knockdown (AHR-KD) cell lines. MAAG and CRP conducted all experiments using endothelial cells. IPS performed the Western blot analyses. CJGP conducted the experiments in colon cells and performed all qPCR analyses. MAAG, PMA, and CNS wrote the manuscript. All authors discussed the data, provided critical feedback, and reviewed the manuscript.

### Notes

The authors declare no competing financial interest.

## ACKNOWLEDGMENTS

This work was supported by EU Horizon 2020 research & innovation programme (H2020-NMBP-BIO-2017) (grant number 760891), European Commission program Teaming for Excellence, Grant No. 101060346, the European Union’s Horizon 2020 research and innovation programme under grant agreement No 951921, NORTE2030-FEDER-01777300 - SCALE-ImmunoHUB2030 supported by Norte Portugal Regional Operational Programme (NORTE 2030), under the PORTUGAL 2030 Partnership Agreement, through the European Regional Development Fund (FEDER); Ludwig Institute for Cancer Research Core Award; COMPETE2030-FEDER-00693400, supported by FEDER and national Funds (Fundação para a Ciência e Tecnologia, FCT), operation number 15824 relative to applications to MPr-2023-12.

iNOVA4Health (LISBOA-01-0145-FEDER-007344; UIDB/04462/2020), co-funded by FCT/Ministério da Ciência e do Ensino Superior (MCTES), through national funds, and by FEDER under the PT2020 Partnership Agreement, is acknowledged. CJGP (grant No. 2022.11465.BD), CRP (grant No. 2023.00453.BD), and IPS (grant No. 2023.02417.BD) are recipients of PhD fellowships from FCT.

PMA, ATJ, RS, YL, and PL acknowledge support from the Ludwig Institute for Cancer Research Core Award. PMA and SM thank funding from the European Union’s Horizon 2020 research and innovation programme under grant agreement No. 951921, NORTE2030-FEDER-01777300 - SCALE-ImmunoHUB2030, and COMPETE2030-FEDER-00693400.

## ABBREVIATIONS USED

AHR: Aryl Hydrocarbon Receptor
AHRR: Aryl Hydrocarbon Receptor Repressor
CH: CH-223191
CVD: Cardiovascular Disease
CYP1A1: Cytochrome P450 Family 1 Subfamily A Member 1
CYP1B1: Cytochrome P450 Family 1 Subfamily B Member 1
DHLC: 11β,13-Dihydrolactucin
DHLCP: 11β,13-Dihydrolactucopicrin
EROD: Ethoxyresorufin-O-deethylase
IBD: Inflammatory Bowel Disease
ICAM-1: Intercellular Adhesion Molecule-1
IFD: Induced Fit Docking
IL-1β: Interleukin-1 Beta
IL-6: Interleukin-6
IL-8: Interleukin-8
KD: Knockdown
LC: Lactucin
LCP: Lactucopicrin
NF-kB: Nuclear Factor Kappa-light-chain-enhancer of Activated B cells
NLRP3: NLR Family Pyrin Domain Containing 3
STL: Sesquiterpene Lactone
TCDD, 2,3,7,8-: Tetrachlorodibenzo-p-dioxin
TIPARP: TCDD-Inducible Poly(ADP-ribose) Polymerase
TNFα: Tumor Necrosis Factor.

